# Metagenomic Analysis of Bacterial Community Associated with Rhizosphere and Phyllosphere of Basmati Rice

**DOI:** 10.1101/2020.04.09.034009

**Authors:** Maria Rasul, Sumera Yasmin, Sughra Hakim, Ahmad Zaheer, Babur Mirza, M Sajjad Mirza

## Abstract

Rice is one of the most important crops for feeding about more than half of the world’s population. Yield of rice crop is significantly hampered by various biotic and abiotic factors. Application of cost-effective and environment-friendly bioinoculant is a common practice to combat the yield losses of rice in this era. The rhizosphere and phyllosphere of rice plants provide specific habitats for various micro-organisms. In the present study, the bacterial population dwelling the rhizosphere and phyllosphere of Basmati rice were explored using metagenomic approach from rice growing areas of Punjab, Pakistan. The bacterial communities associated with the rice rhizosphere of different rice growing areas as well as phyllosphere/rhizosphere were compared. Out of 20,069 16S rRNA gene sequences retreived from rhizosphere soil, 6485 were originated from Faisalabad, 5174 from Gujranwala and 8410 from Sheikhupura. Data analyses revelead that *Proteobactria* was dominant phylum at all three sites. *Choloflexi* was second abundant phylum at Sheikhupura, while *Actinobacteria* and *Firmicutes* at Gujranwala and Faisalabad. In addition to dominant culturable PGPB *Bacillus* genus at all three sites, *Nitrosospira, Gaiella, Marmoricola, Clostridium sensu stricto*. Maximum geners were detected from Faisalabad (159), followed by Sheikhupura (146) and Gujranwala (131). Comparison of the common sequences at the genus-level revealed that maximum number of shared genera (101) were observed in Sheikhupura and Gujranwala. 50 genera were specifically related to Faisalabad, while 27 and 21 genera were detected for Sheikhupura and Gujranwala.

In the phyllosphere, *Proteobacteria* (79.6%) was detected as dominant phylum followed by *Firmicutes* (9.8%), *Bacteroidetes* (8.6%), *Chloroflexi* (4.3%) and *Actinobacteria* (0.9%). Comparison of phyllosphere and rhizosphere showed less bacterial diversity in the phyllosphere but *Bradyrhizobium, Sphingomonas*, GP6, *Pseudomonas, Bacillus* are abuundant. Fifteen genera were detected at both compartments. Furthermore, we may select these strains for development of compatible inocula for application in the rhizosphere and phyllosphere of rice.

## Introduction

Rice production depends on chemical fertilizers and pesticides. Excessive use of chemical fertilizers disturb the balance of macro and micronutrients in soils. Diverse microbial population colonize different parts of the rice plant. The diversity and activity of microbial communities affect the soil fertility and the nutrient use efficiency [1]. In recent years, research on the diversity of plant-associated microbiota “plant microbiome” has made considerable progress. The analysis of the plant microbiome includes linking microbial ecology with the biology and functioning of the host plant. It also helps to explore these microorganisms as a reservoir of additional genes and functions for their host [2].

The rhizosphere is an area (<2 mm) around the roots. It is specific microbial habitat in the soil ecosystem and a hot-spot for many organisms [3]. Rhizospheric microbial communities comprised of algae, bacteria, fungi, viruses, oomycetes, arthropods, nematodes, archaea and protozoa [4,5]. Among these microbes, most of the members are part of intricate food chain which utilize the minerals nutrients secreted from plants. In this area, the soil is swayed by roots by deposition of plant-exudates. In addition, rhizosphere and root microbiota provide beneficial effects on the plant i.e. improved nutrients uptake from soils for plant growth and protection against pathogens [6]. Rhizosphere organisms have been characterized into three different groups. Members of first group include P-solubilizer, nitrogen fixers, biocontrol agents, mycorrhizal-fungi and protozoa and have beneficial effect on crop health. It. Members of second group includes bacteria, pathogenic fungi, oomycetes and nematodes that have deleterious effects on crop growth. Members of third group are the human pathogens. During the last decade, the spread of human pathogenic bacteria residing inside plant tissues and on the surface has been frequently reported [7]. Knowledge about processes that influence and drive the dynamics and composition of the soil microbiota remains necessary to secure the human health along with plant productivity. Different members of these microbial communities commonly interact with one another and with their hosts. Therefore, it is mandatory step to grasp the microbial diversity as much as possible. Furthermore, understanding of microbial community structure is also essential to explore the individual function of bacteria, fungus, etc.

Rhizospheric abundance/ richness of microbial species has been suggested as an indicator of the diversity and yield of plants above ground under various environmental conditions [8,9]. Previously, cultivation-based technique is predominantly used to assess the microbial diversity in rhizosphere. However, the the major limitation of this method for plant-biologists and microbiologists is that only 1% of microbial community members could be cultivated under controlled laboratory through this technique, while remaining 99% of microorganisms inhabiting the rhizosphere remain unexplored [10]. Nevertheless, ecologists are facing a challenge to correlate the microbial diversity present in the rhizospheric soil with their function in the natural ecosystem.

The modern high-throughput sequencing technique enables us to assess and explore the microbiota from the challenging environments i.e. oceans, hot springs, etc. through culture-independent method (metagenomic) is well documented [11,12]. Metagenomics the “modern analytical tools” is a fast-growing field of genome sciences, that divulge the functional potential of the rhizosphere microbiota. This technique or tool explained the characteristics of intact microbes in their natural habitats and their interactions with host plant, for understanding different intrinsic processes like nutrient cycling, ecosystem function and carbon sequestration [11,12].

In addition to studies of rhizosphere-plant microbiome, few reports are available revealing the substantial abundance and diversity of microbes present on aerial part of plant known as phyllosphere [13]. These diverse microbial community present on phyllosphere include bacteria, fungi, viruses, algae, archaea and rarely nematodes and protozoa [14]. Bacteria surpass in abundance and diversity by other detected groups [15]. Bacteria residing on the phyllosphere of rice and their physiological adaptations to the habitat have not been explored as much as the rhizosphere residing bacteria were studied. The reason behind is that aerial parts of plant are inhospitable, open system for microbes that are highly swayed by variable environmental conditions and nutrient deficiency. Therefore, microbial species manage to colonize the phyllosphere using different mechanisms. To date, a number of bacteria from the phyllosphere of rice have been isolated and characterized for the beneficial potential like nutrient acquisition, phytohormone production or protection against pathogens [16-18].

Previously, research work was conducted to study the microbial community residing as root endophytes of Indian rice [19], rhizospheric microbiota of two Venezuelan rice varieties [20] and phyllosphere analysis in two major rice growing areas of China and the Philippines [21]. According to our knowledge, no microbiome information is available from aromatic Basmati rice growing areas. It is first report which targeted the unexplored aromatic Basmati rice growing areas of Pakistan for rhizosphere microbial diversity. Keeping in view the effect of rhizosphere and phyllosphere microbiome on growth promotion and suppression of foliar pathogen of rice, the present metagenomic study was conducted to investigate the microbial population dwelling the rice rhizosphere and phyllosphere for development of compatible inoculum. We focused to identify the dominant bacterial genera colonizing the rice: (1) differences in overall rhizosphere microbiota of Basmati rice in different fields of rice “Kalar belt”, (2) diverse bacterial community structure of phyllosphere and rhizosphere.

## Materials and Methods

### Soil Sample Collection

Rhizospheric soil samples were collected from rice plants growing in the experimental areas of NIBGE field, Faisalabad (31°23’44.0"N 73°01’37.3"E), Adaptive research station Sheikhupura (31°42’42.6"N 73°59’07.5"E) and Adaptive research station Gujranwala (32°13’21.5"N 74°13’02.7"E) Gujranwala for metagenomics analysis. For rhizospheric soil sampling, ten plants were uprooted from different fields of each site. Loosely attached soil from all plants was removed and root adhered soil <2 mm were collected from all roots and homogenized to make a composite sample.

### DNA Extraction from Rhizosphere Soil and Phyllosphere of rice by Culture-Independent Approach

For genomic DNA extraction from rice rhizosphere soil, Bead beater was used for mechanical lysis and DNA isolation kit (MP Biomedicals, Santa Ana, California, USA) was used following the instructions as described by manufacturer. “Lysing matrix E” in 2 mL tube was used for rhizosphere soil samples, that had a glass bead (4 mm), ceramic spheres (1.4 mm) and silica spheres (0.1 mm) [22].

For genomic DNA extraction from rice phyllosphere, ten leaves per plants were taken from each plant and considered as an individual sample. Bead beater was used for mechanical lysis and FastDNA SPIN kit (MP Biomedicals, Santa Ana, California, USA) was used following the manufacturer instructions. “Lysing matrix A” was used for leaf samples, which contain garnet matrix and 1/4 ceramic spheres in 2 mL tube. For homogenization of bacterial cells with lysis matrix A “CLS-TC” cell lysis solution was used [23,24].

### Metabarcoding of 16S rRNA Gene and Barcoded Illumina Sequencing

16S rRNA gene was amplified from the DNA, isolated from rhizosphere soil and phyllosphere using two steps PCR approach. First step, consisted of one set of primer including 16S rRNA gene primer and Illumina sequencing primer. In the second step of PCR multiple primers were used including Illumina adapter and sequencing-primer sequence (Table 1). MiSeq. platform (Illumina) was used for pair end sequencing from both forward and reverse direction. Fastq files of 16S rRNA gene sequences were submitted to NCBI Sequence Read Archive (SRA) under BioProject ID <u>PRJNA574892</u> (https://www.ncbi.nlm.nih.gov/bioproject/574892).

**Table 1.**
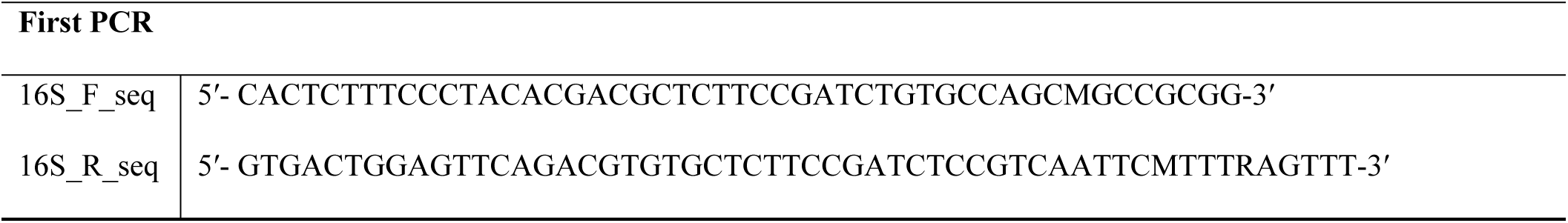
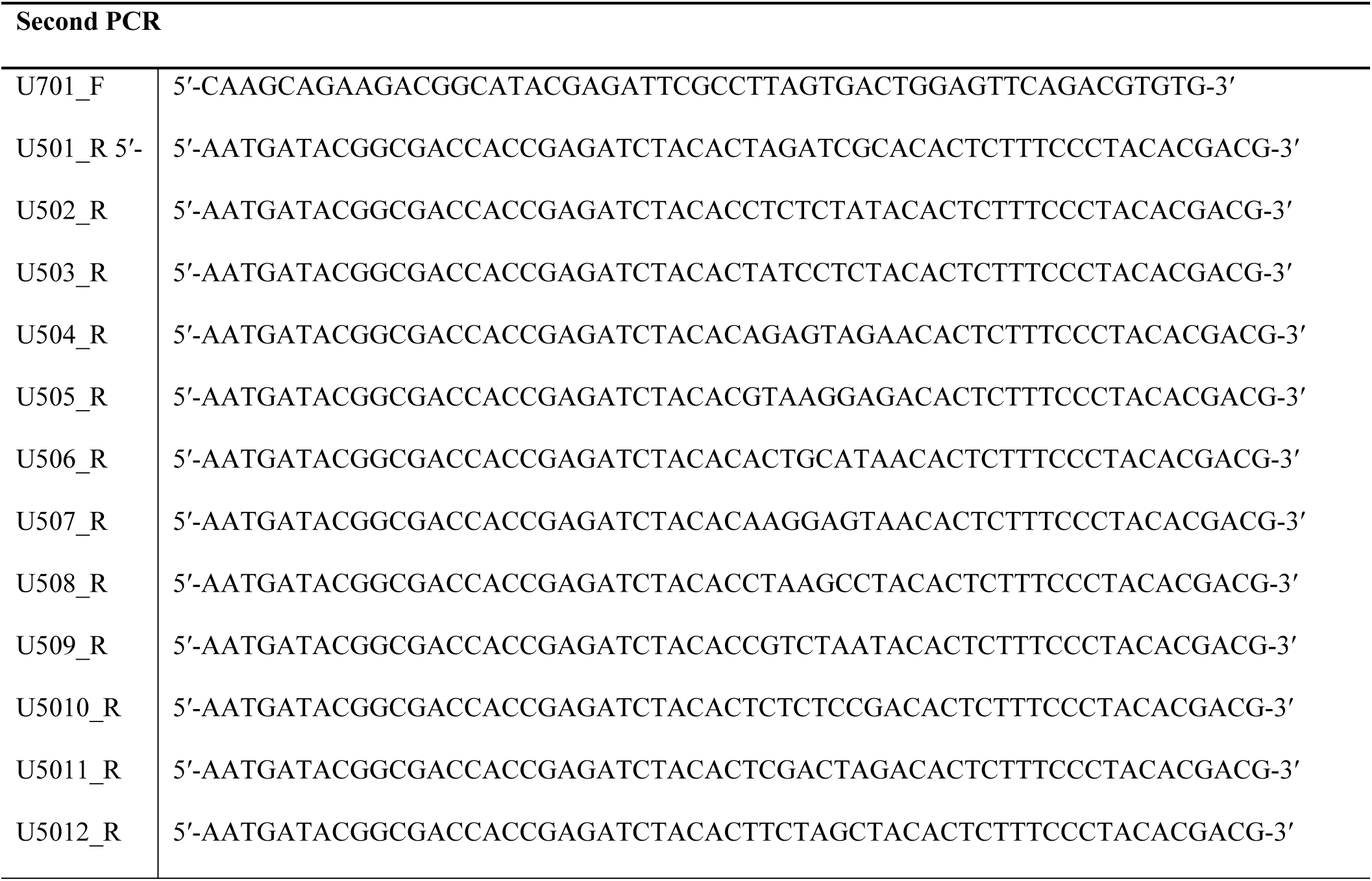
List of Primers Used for PCR Amplification and Sequencing.

### 16S rRNA Gene Sequence Analysis

16S rRNA amplified sequences from phyllosphere and rhizosphere soil were initially processed for quality control parameters. Mothur software (Package v.1.35.1) was used to assemble the Paired-end Illumina reads. Quality of the sequences were improved by trimming and filtering [25,26]. Terminal parts of sequences containing primer sequences were trimmed and the sequences in which forward and reverse are not aligned were filtered out. A maximum expected error of 0.5 was used and the sequences were cut to a minimum of 370 bp read length, maximum eight base homopolymer, sequences with unidentified bases (N) and sequence >1 incorrect match with unique bar code identifier. Mothur was used for identification and removal of chimeric sequences. High quality sequences from phyllosphere and rhizospheric soil were identified and processed through Ribosomal Database Project (RDP; http://rdp.cme.msu.edu) and Naive Bayesian Classifier 2.5 [27]. These sequences were classified, aligned and clustered (at 97% DNA similarity) into operational taxonomic units (OTUs). Singleton were eliminated to reduce the sequencing error. Rarefaction curves were generated using 97% similarity from triplicate samples of each experimental site. Richness (Chao1), diversity (Shannon index) and evenness were calculated at 97% similarity. All sequences have been deposited in the GenBank Sequence Read Archive (SRA) for accession numbers.

## Results

### Bacterial Diversity Analysis Based on 16S rRNA Gene in the Rhizosphere of Basmati Rice at Different Sites by using Illumina Sequencing

Bacterial DNA was extracted from the rice rhizospheric soil. Bacterial diversity present in rhizosphere of three rice growing areas of Pakistan were explored by 16S rRNA gene-based sequence analysis. Overall 20,069 16S rRNA gene sequences were retrieved from the rice rhizosphere soil that was collected from Basmati rice growing areas. Out of the total retreived sequences, 6485 sequences were originated from Faisalabad 5174 from Gujranwala soil and 8410 from Sheikhupura field soil.

Rarefaction curves based on OTUs at 3% dissimilarity level were used to determine the sequencing depth. Figure 1 rarefaction curves depicted the bacterial diversity and richness of the samples. Maximum genetic diversity was captured from the soil sample collected from all three site as indicated by plateaue curve.

**Figure 1.**
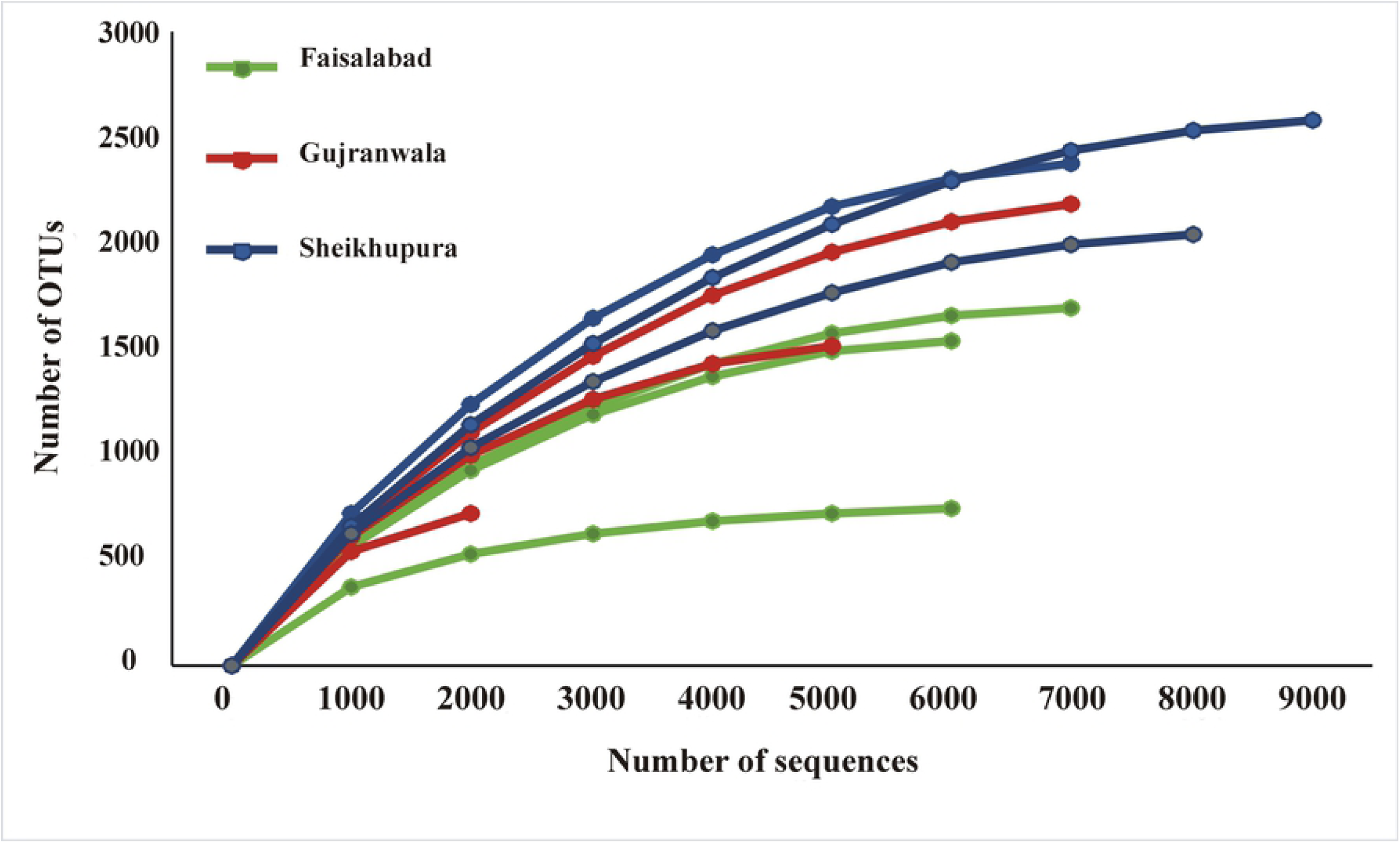
Rarefaction Analysis of Rhizospheric Soil Sample of Basmati Rice Collected from Basmati Rice Growing Areas of Kalar Belt and Faisalabad. Rarefaction Curves at 97% Sequence Identity across Different Samples

Diversity indecies were used to quantify the biodiversity in terms of number of species as well as its abundance in particular habitat. Bacterial richness (2339±226) and shannon diversity index (H= 7.01±0.17) observed for Sheikhupura site were significantly higher as compared to those for Gujranwala (H= 6.59±0.41) and Faisalabad (6.27±0.44) field. Evenness in rhizosphere soil samples collected from Gujranwala (E= 0.91±0.015) was higher as compared to those of Sheikhupura (E=0.90±0.017) and Faisalabad (E= 0.88±0.007) (Table 2).

**Table 1.** Evenness, Diversity and Richness of Rhizospheric Soil Sample of Basmati Rice at 3% Sequence Divergence.

| Sites | N | Clusters | Chao | H' | varH | E |
| --- | --- | --- | --- | --- | --- | --- |
| **Faisalabad** | 6485 ± 485 | 1327 ± 509 | 1330 ± 507 | 6.27 ± 0.44 | 0.0004 | 0.88 ± 0.007 |
| **Gujranwala** | 5174 ± 2904 | 1488 ± 744 | 1489 ± 743 | 6.59 ± 0.41 | 0.0005 | 0.91 ± 0.015 |
| **Sheikhupura** | 8410 ± 921 | 2338 ± 226 | 2339 ± 226 | 7.01 ± 0.17 | 0.0003 | 0.90 ± 0.017 |

± showed standard deviation of three replicates. Alpha diversity within each community using two commonly used metrics. Richness (chao) measures how many unique OTUs were detected in each sample while Shannon-entropy measures the randomness or uncertainty in community. E, Shannon Wiener equitability index; H', Shannon-Wiener index; varH', variance of H';

### Global Beta Diversity of Different Soils Collected from Rice Kalar Belt

Beta diversity indices were use for quantitative estimation of differnces in divesity between three soils. Minimum beta diversity was observed in soil of Sheikhupura followed by Faiasalabad and Gujranwala. Values for diversity analysis ranged from 0.39-0.52 for Whittakar, 0.19-0.26 for Harrison, 51-60 for cody, 0.10-0.13 for Routledge, 0.48-0.64 for Wilson-shmida, 0.23-0.32 for Mourelle, 0.12-0.16 for Harrison 2 and 0.19-0.24 for Williams. Values of all parameters for beta diversity analysis were lower for Sheikhupura and maximum for Gujranwala except Harrison 2 and Williams (Table 3).

**Table 3.** Global Beta Diversities Detected by Illumina Sequencing Based on 16S rRNA Gene in the Rhizospheric Soil Samples of Basmati Rice.

| Parameters | Sites of Basmati Growing areas |  |  |
| --- | --- | --- | --- |
|  | Faisalabad | Gujranwala | Sheikhupura |
| **Whittakar** | 0.42 | 0.52 | 0.39 |
| **Harrison** | 0.21 | 0.26 | 0.19 |
| **Cody** | 60 | 56 | 51 |
| **Routledge** | 0.11 | 0.13 | 0.10 |
| **Wilson-Shmida** | 0.53 | 0.64 | 0.48 |
| **Mourelle** | 0.27 | 0.32 | 0.23 |
| **Harrison 2** | 0.16 | 0.12 | 0.13 |
| **Williams** | 0.24 | 0.19 | 0.21 |

All values were an average of three replicates

Sequence analyses indicated that a significant fraction (58-70%) of total retrieved sequences from three sites were belonged to dominant taxonomic groups like *Proteobactria, Acidobacteria, Actinobacteria, Choloflexi* and *Firmicutes. Proteobacteria* was the dominant phyla at all three sites comprising 24% of total retrieved sequences from Faisalabad, 22% from Sheikhupura, 19% from Gujranwala. *Firmicutes* were (17%) *Chloroflexi* (18%) and *Actinobacteria* (12%) were dominant phyla after *Proteobacteria* in Faisalabad, Sheikhupura and Gujranwala, respectively. Comparison of these retrieved sequences among these sites revealed that “*Chloroflexi*” at the Gujranwala and Sheikhupura were relatively under representative like 10% and 7.6% as compared to Faisalabad site 18%. Whereas *Actinobacteria, Acidobacteria* and *Fermicutes* were abundant at Faisalabad and Gujranwala as compared to Sheikhupura. A significant fraction of total retrieved sequences from all three sites were uncultured comprising of 20-32%. Minor OUTs (presenting ≤ 1%) formed 8-10% of total sequences across all tested sites (Table S1 and Figure 2). Component analysis (CA) showed that Component 1 and Component 2 contributed about 99% of total variance. (Figure 3). Figure showed that most of the bacterial genera in all three sites are similar except few ones.

**Figure 1.**
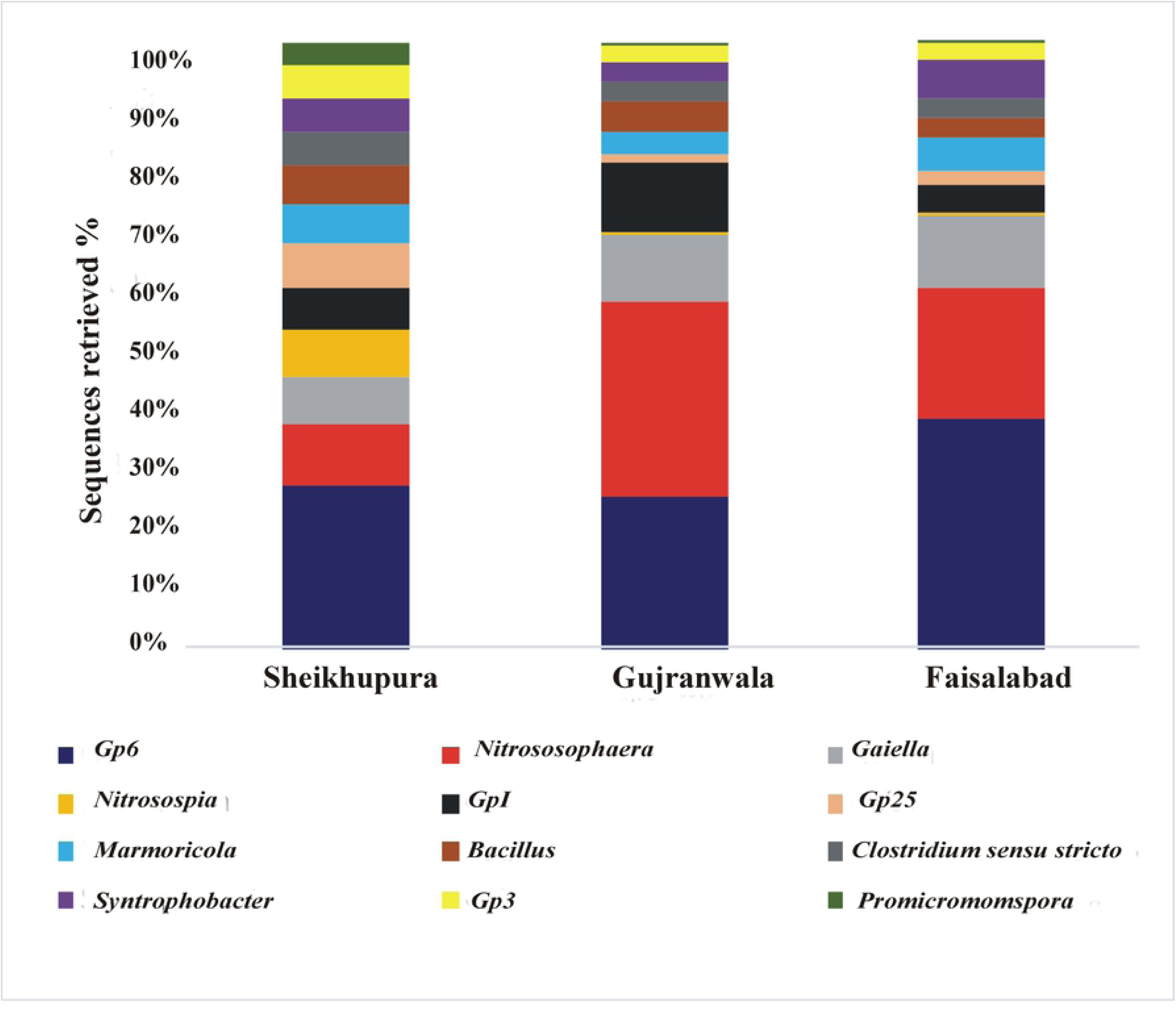
Relative Abundance of Dominant Bacterial-Phyla Detected by Illumina Sequencing of 16S rRNA Gene in the Rhizosphere Soil Sample of Basmati Rice Collected from Different Areas of Kalar Belt and Faisalabad.

**Figure 3.**
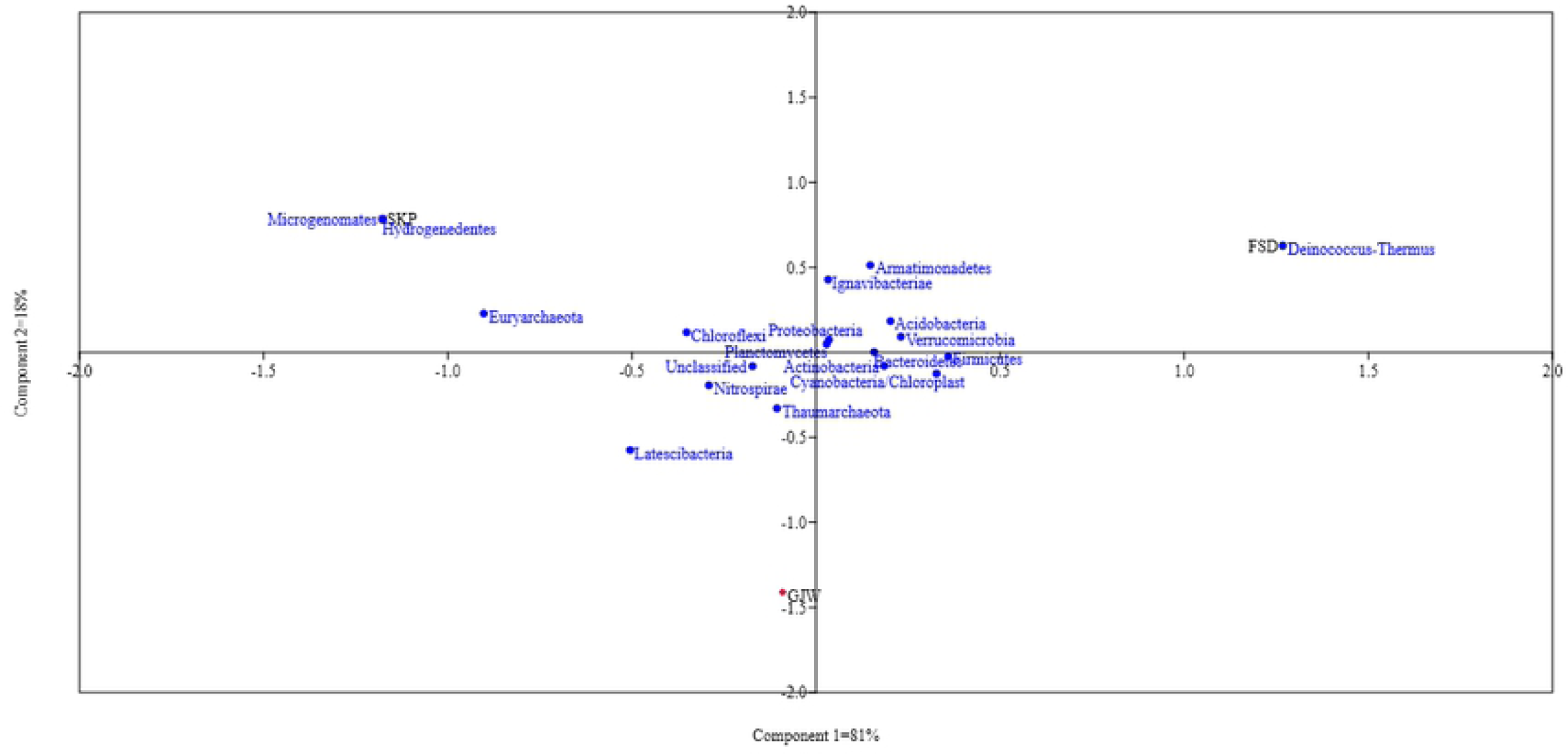
Coordinate Analysis (CA) Profile of Microbial Diversity Across All Rhizosphere Samples using Bray-Curtis Dissimilarities.

### Relative Distribution of Bacteria in Different Rhizosphere Samples at Genus Level

Gujranwala, Faisalabad and Sheikhupura are three sites of aromatic Basmati rice growing Kalar belt. The results of classification at genus level for each soil sample collected from these three sites were depicted in Figure 4, Table S2. GP6, *Nitrososphaera* and *Gaiella* genus contributed more than 1%. A total of 159 geners were detected from rhizosphere soil of Faisalabad, 131 from Gujranwala and 146 from Sheikhupura. Comparison of the shared sequences at the genus level revealed that maximum number of shared genera (101) were observed in Sheikhupura and Gujranwala. 50 genera were specifically to rhizosphere soil of Faisalabad, while 27 and 21 genera were specifically detected for Sheikhupura and Gujranwala, respectively Figure 5.

**Figure 4.**
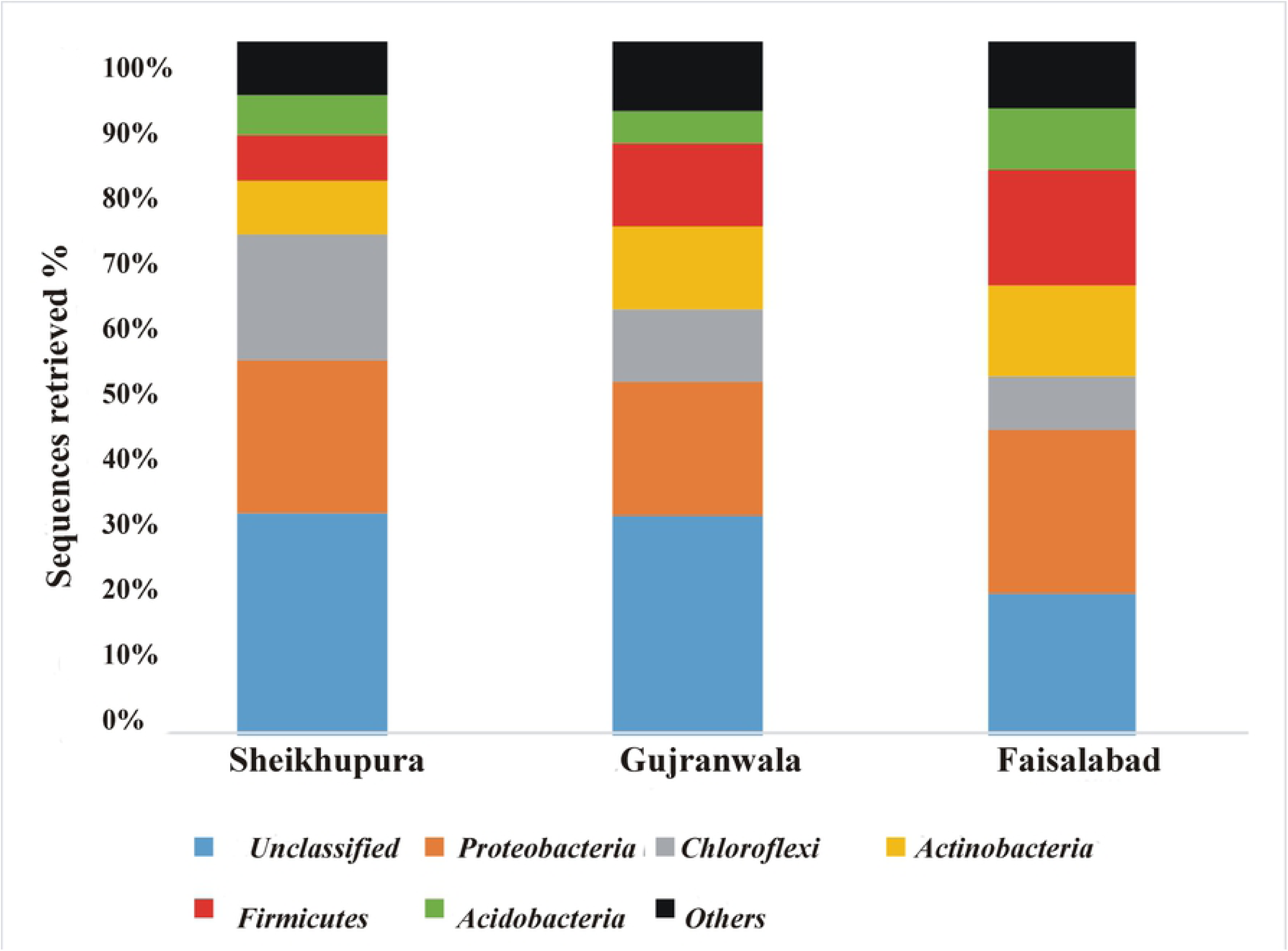
Relative Abundance of Dominant Bacterial-Genera Detected by Illumina Sequencing of 16S rRNA Gene in the Rhizosphere Soil Sample of Basmati Rice Collected from Different Areas of Kalar Belt and Faisalabad.

**Figure 5.**
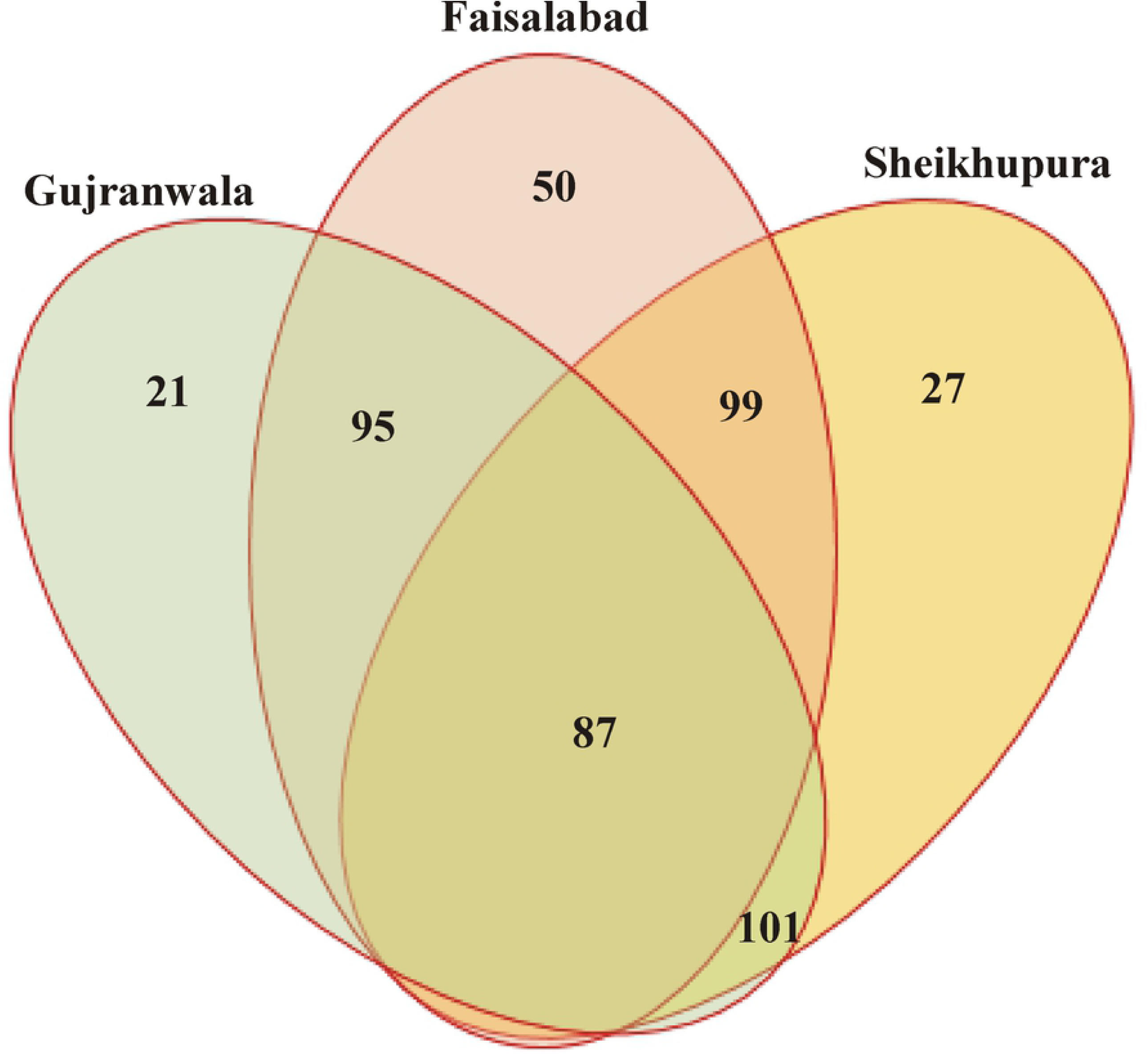
Venn Diagram Showing the Bacterial Genera Shared among the Rhizosphere Soil Samples of Basmati Rice Collected from Different Areas of Kalar Belt and Faisalabad.

### Compartive Abundance of Plant Growth Promoting Genera at Different Soils of Rice Belt

Comparison of retreived sequences from three different soils (Figure 6) revealved that among 227 genera, 15 bacterial genera were reported as potental phosphate solubilizers, 15, 15 were reported as nitrogen fixer and IAA producer. Among phosphate solubilizing bacteria, *Bacillus* was dominant genus followed by *Promicromonospora* and *Agromyces* in the rhizospheric soil sample collected from Faisalabad. *Bacillus* follwed by *Paenibacillus* and *Mycobacterium* were dominent genera in soil collected from Gujranwala while *Bacillus* and *Thiobacillus* were dominent genera detected from soil of Sheikhupura field (Figure 6).

**Figure 6.**
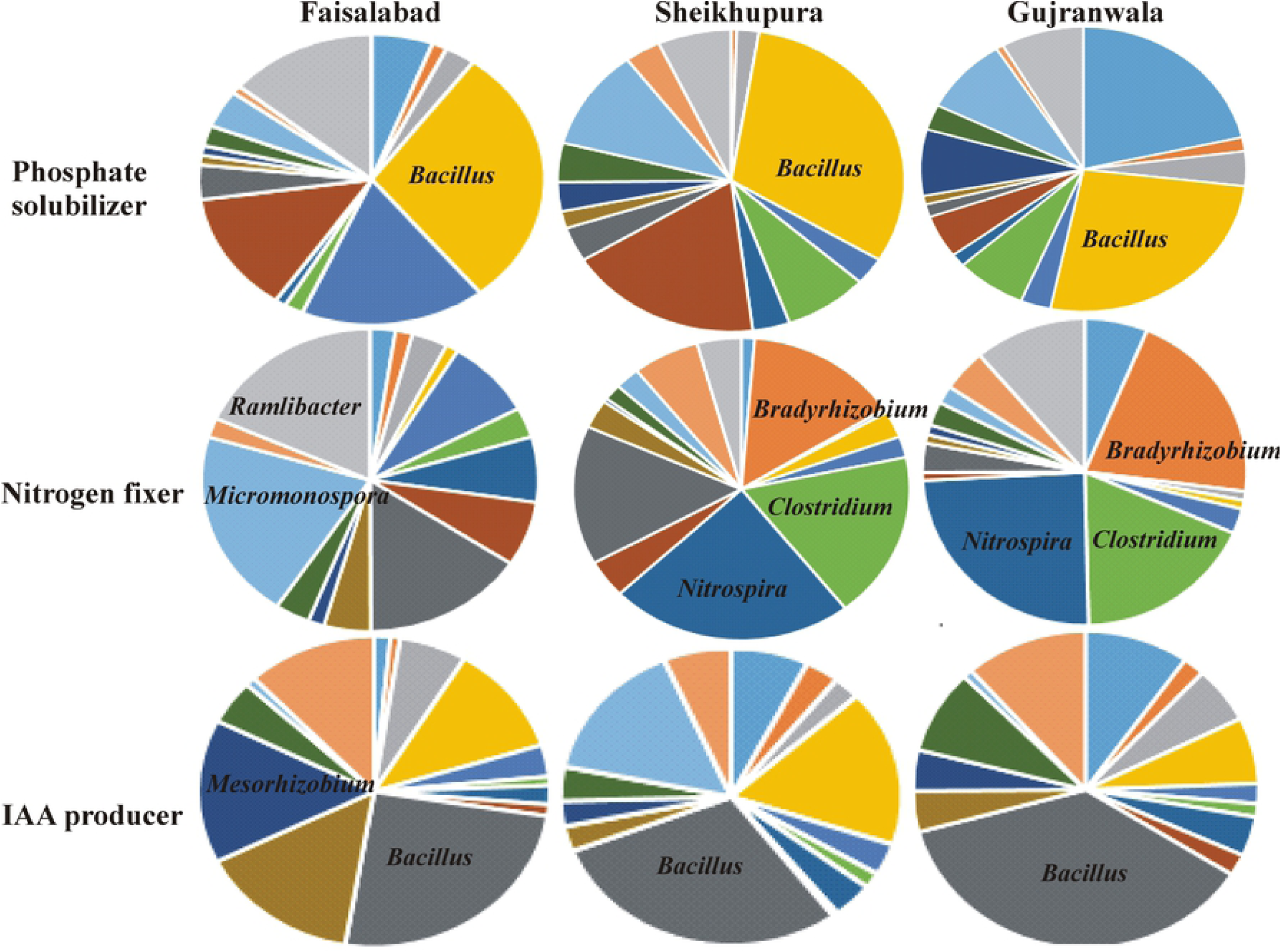
Relative Abundance of Dominant Bacterial-Genera Detected by Illumina Sequencing of 16S rRNA Gene in the Rhizosphere Soil Sample of Basmati Rice Collected from Different Areas of Kalar Belt.

Among nitrogen fixing bacteria *Promicromonospora, Ramlibacter, Paenibacillus* and *Ensifer* were dominently detected genera in soil collected from Faisalabad. *Nitrospira, Clostradium* and *Braydirhizobium* were dominent nitrogen fixing bacterial genera detected from soils colleted from Gujranwala and Sheikhupura field.

Potential genera having IAA producing abiltity were *Bacillus, Mesorhizobium, Promicromonospora, Agromyces, Paenibacillus* and *Pseudonocardia* in soil collected from Faisalabad. *Bacillus, Paenibacillus, Pseudonocardia* and *Streptomyces* were dominant genera in the soil of Gujranwala. *Bacillus, Agromyces, Aeromicrobium* and *Paenibacillus* were dominant in the soil collected from Sheikhupura (Figure 6).

Seven common bacterial genera i.e. *Azospirillum, Bacillus, Brevibacillus, Mesorhizobium, Paenibacillus, Streptomyces* and *Sphingomonas* have potential for IAA production, P-solubilization and nitrogen fixation.

### Comparison of Bacterial Diversity at phylum level in the Rhizosphere and Phyllosphere

For application of any biocontrol agent against rice foliar pathogen i.e. Bacterial leaf blight, we tried to exlore the bacterial community residing in the phyllosphere of rice in comparison of the rhizosphere bacterial community in rice plant of the selected Faisalabad site.

Comparison of bacterial phyla in the rhizosphere and phyllosphere showed abundance in the rhizospheric soil as compared to phyllosphere. A total of 9383 16S rRNA sequences were retrieved from rhizospheric soil and 56714 sequences (including chloroplast sequences) were retrieved from phyllosphere. For further analysis choloplast sequences were removed from total retreived sequences of phyllosphere. Diversity indecies were used to quantify the biodiversity in terms of number of species as well as its abundance in particular habitat. Bacterial richness (20350±2800), shannon diversity index (H= 9.55±0.086) and evenness in rhizosphere soil samples collected from rhizosphere (E=0.928±0.009) was higher as compared to those of phyllosphere (E=0.3145±0.028) of rice plant (Table 4).

**Table 4.** Evenness, Diversity and Richness of Rhizospheric Soil Sample and phyllosphere of Basmati Rice at 3% Sequence Divergence.

**Basmati Rice at 3% Sequence Divergence**
| Compartments | N | clusters | chao | H' | varH | E |
| --- | --- | --- | --- | --- | --- | --- |
| Phyllosphere | 60346±11397 | 4297±645 | 20350±2800 | 2.63±0.23 | 0.000177 | 0.3145±0.028 |
| Rhizosphere | 55658±13895 | 29874±5922 | 82310±16789 | 9.55±0.086 | 5.67E-05 | 0.928±0.009 |

± showed standard deviation of three replicates. Alpha diversity within each community using two commonly used metrics. Richness (chao) measures how many unique OTUs were detected in each sample while Shannon-entropy measures the randomness or uncertainty in community. E, Shannon Wiener equitability index; H', Shannon-Wiener index; varH', variance of H';

Beta diversity indices were use for quantitative estimation of differnces in divesity between two compartments. Beta diversity decreases with increase in sample number. The maximum decrease numbers of specimens consistently found with the rhizosphere, Whittaker- 0.49, Harrison- 0.25, Routledge 0.13, Wilson-shmida- 0.6, Mourelle 0.3, Harrison-2 0.16 ans Williams 0.245 as compared to phyllosphere, Whittaker- 1.68, Harrison- 0.83, Routledge 0.41, Wilson-shmida 1.71, Mourelle 0.86, Harrison-2 0.64 ans Williams 0.56 (Table 5).

**Table 5.**
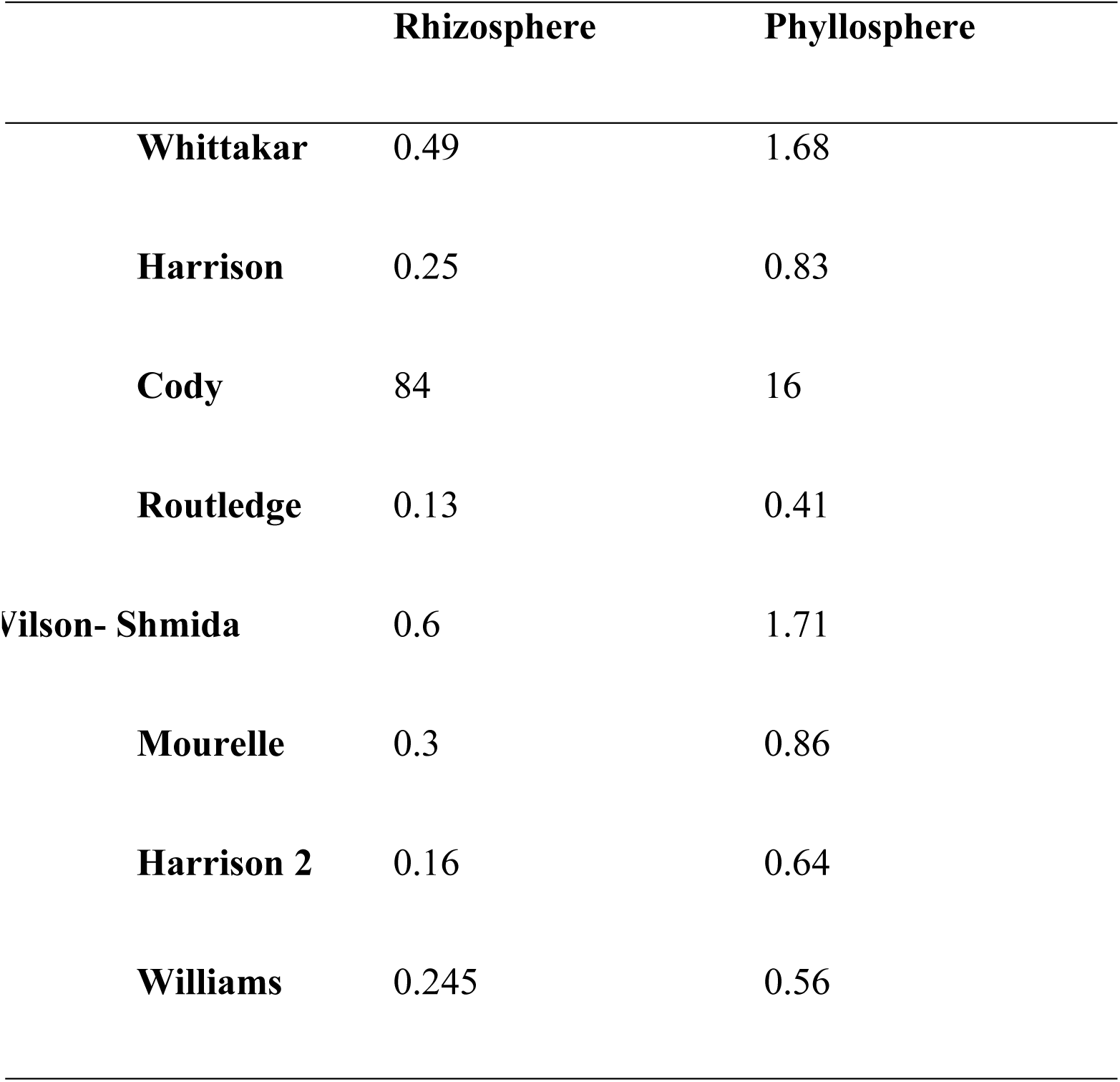
Comparasion of Global Beta Diversities Detected by Illumina Sequencing Based on 16S rRNA Gene in the Rhizospheric Soil and phyllosphere Samples of Basmati Rice.

Sequences retrieved from rhizosphere belonged to 18 different phyla while 7 phyla were detected from phyllosphere sample of rice. Among the detected phyla, *Proteobacteria* was dominant in both compartments i.e. phyllosphere (79.6%) and rhizosphere (37.2%). Total sequences retrieved from phyllosphere showed significant fraction of *Proteobacteria* (79.6%) followed by *Firmicutes* (9.8%), *Bacteroidetes* (8.6%), *Chloroflexi* (4.3%), *Actinobacteria* (0.9%) (Table S3). Seven bacterial phyla were detected from both compartments i.e rhizo and phyllosphere (Figure 7).

**Figure 7.**
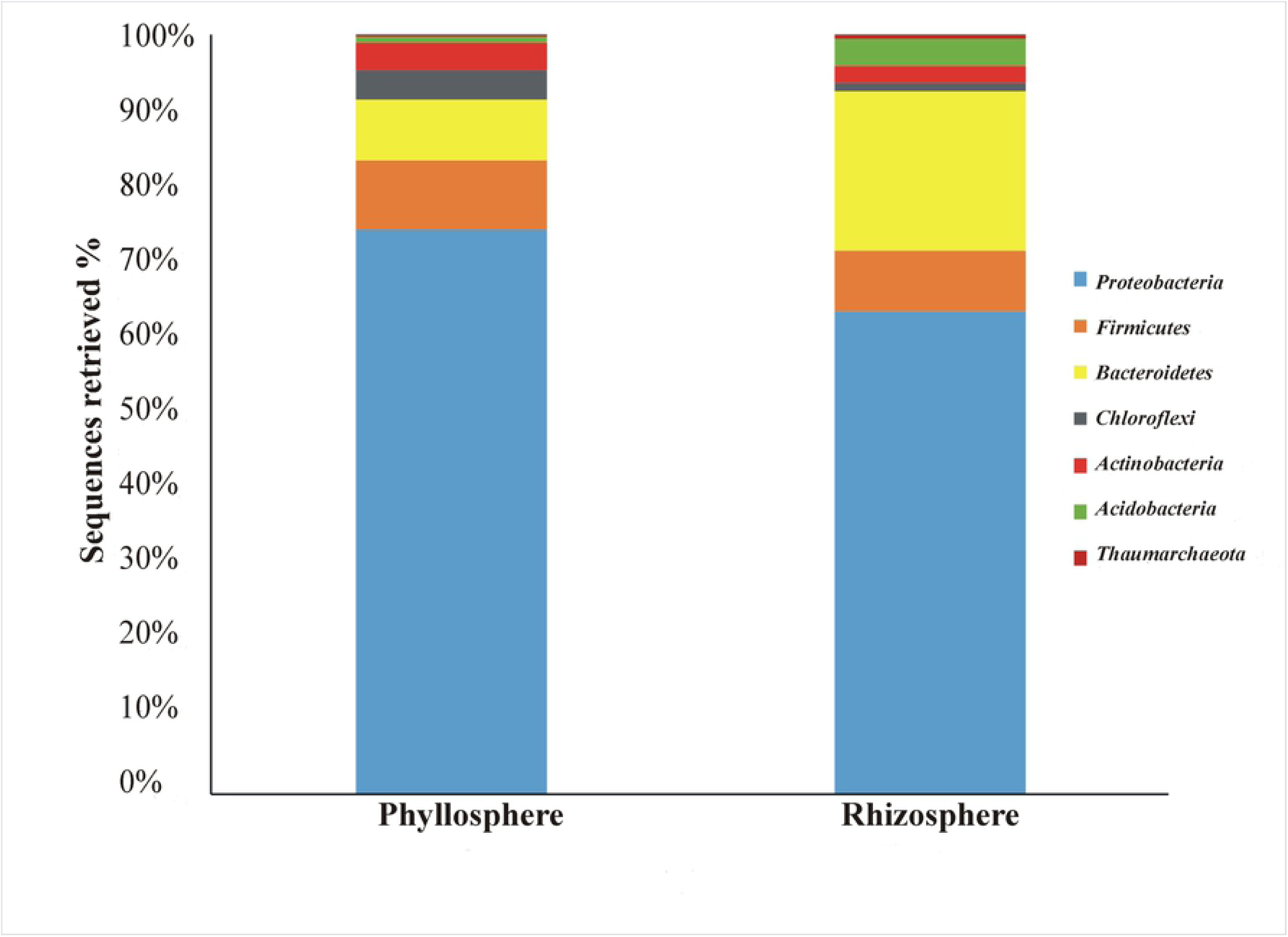
Relative Abundance of Dominant Bacterial-Phyla Detected by Illumina Sequencing of 16S rRNA Gene in the Phyllosphere and Rhizosphere of Basmati Rice.

### Relative Distribution of Bacterial Genera within Rhizosphere and Phyllosphere of Basmati Rice

16S rRNA based sequences analyses from phyllosphere and rhizosphere revealed the diversity at genus level. A total of 208 different genera were detected in rhizospheric soil sample (Table S4), while only 24 genera were detected from the phyllosphere (Table S5). From phyllosphere, *Bacillariophyta* (22%) was the dominant genus, followed by *Sphingomonas* (9%) and *Bradyrhizobium* (7%). While a significant fraction i.e. 81% of total retrieved sequences from the rhizospheric soil samples could not be classified at genus level. *Thaurea* was the dominant genus in rhizosphere contributing 3.6% of total sequences followed by *GpI* 1.5%, *Ohtaekwangia* 1.18% and Gp6 1.17%. Other 204 genera have presented by <1% and contributed 11.3% in total retrieved sequences. In the comparative analysis 15 genera were common in the bacterial community of phyllosphare and rhizosphere. These 15 genera represented 52% of phyllosphere sequences and 3.7% of rhizospheric soil sample (Figure 8).

**Figure 8.**
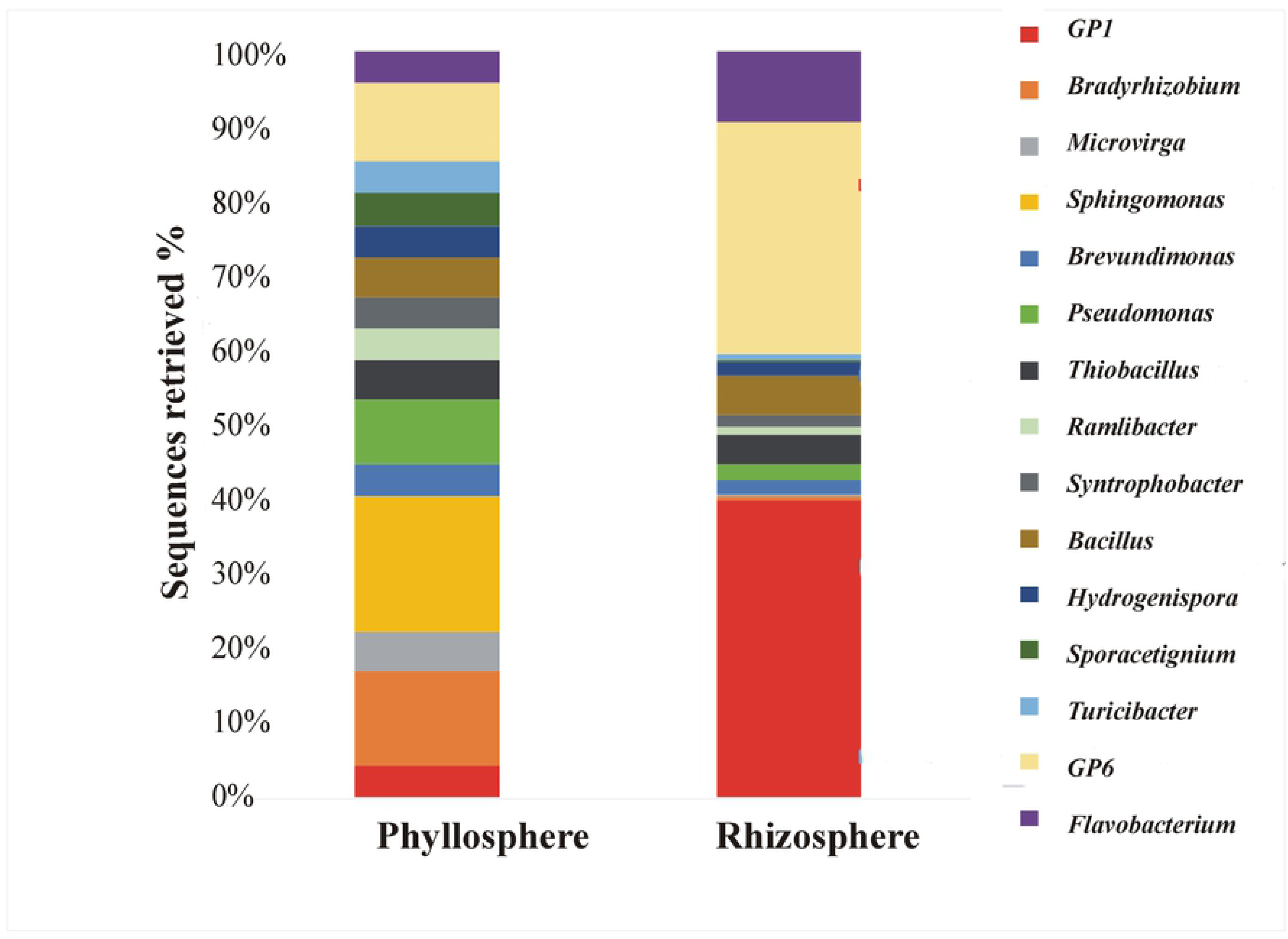
Relative Abundance of Dominant Bacterial Genera Detected by IIlumina Sequencing of 16S rRNA Gene in the Phyllosphere and Rhizosphere of Basmati Rice.

## Discussion

Rice is a highly valued cash crop, important agriculture product and holding special importance as a global staple food. It is a kharif season crop and this cropping season is appropriate for cultivation of Basmati (aromatic) and non-Basmati rice. The premium quality aromatic Basmati rice are grown along with the ‘Kalar belt’ that have clay-soil texture with good water holding capacity [28]. Astonishing number of microbes colonized the rice plants and their abundance is found very high as compared to number of plant cells [7]. Aerial parts (phyllosphere) and the root adhered soils (rhizosphere) of rice plant are important habitats for diverse microbial population [29]. The phyllosphere is dominated by the leaves and severely affected by plant pathogens due to direct contact with fluctuating environment. Therefore, study of microbial community inhabiting the leaf tissues is important for development of bacteria-based biopesticides against foliar pathogen like *Xanthomonas oryzae* causing bacterial leaf blight in rice.

The rhizosphere is a biologically active zone enriched with microbes. The chemical, physical and biological characteristics of this particular zone have a great impact on the plant roots. The soil microflora has a great contribution in nutrient cycling. Therefore, application of the potential beneficial microbes is highly applicable to enhance crop production as well as soil fertility. Actually, these microbes increased the level of essential macronutrients (N, P) for plant uptake. Among microbial communities, the bacterial microbiome particularly has gain much more attention for sustainable agriculture as compared to other groups [30].

In the present study, first time we targeted the unexplored locations of Basmati rice growing Kalar belt and Faisalabad to discern the rhizospheric bacterial diversity. We have observed significant differences in growth and grain yield of rice variety Super Basmati grown in these soils. We hypothesized that the abundance, diversity and distribution of microflora have huge impact of environmental factors, including soil type and crop variety. There is also difference in microflora of soils present along the Kalar belt as compared to Faisalabad that is out of this premium zone of Kalar belt. Secondly, we compared the rhizosphere and phyllosphere bacterial diversity of rice plant grown under natural field conditions. Previous reports revealed that among the different groups of microorganism, bacteria is predominant group present in the rhizosphere of plant [31]. Keeping in view the bacterial abundance, in the current study, Illumina sequencing based on 16S rRNA gene from DNA of rhizospheric soil samples were conducted for estimating the bacterial diversity across three investigated sites of Kalar belt. Diversity indices compares dominance of bacterial species, relative abundance, species richness, evenness. Similarity and dissimilarity related coefficients as measures of alpha and beta diversity is a common practice. Alpha diversity increases with increase in sample number while beta diversity decreases with increase in sample number [32]. Dominant taxonomic groups were *Proteobacteria, Acidobacteria, Actinobacteria, Choloflexi*, and *Firmicutes* in all soil samples. Dominance of *Actinobacteria* and *Proteobacteria* has also been documented in the rhizospheric soil of maize [33].

Previous reports have shown that *Proteobacteria, Acidobacteria, Actinobacteria, Bacteroidetes* and *Fermicutes* were dominant phyla in the rhizosphere soil of *Arabidopsis* and cotton [34-36]. In case of rice, *Proteobacteria, Acidobacteria, Actinobacteria, Fermicutes* and *Planctomycetes* were found to be dominant bacteria among all bacterial communities [37]. *Proteobacteria* remains the most abundant soil bacterial community. *Actinobacteria* are important in the recycling of nutrients [38-40]. *Choloflexi* bacteria are important for decomposition of organic matter [41]. Group of *Firmicutes* and *Proteobacteria* include beneficial bacteria that have potential for siderophores production, IAA production, nitrogen fixation and inhibition of different phytopathogens [42]. It was also found that disease-suppressive soils are rich in bacteria belonging to *Acidobacteria* and *Actinobacteria*. Moreover, these taxa are reported to supress disease-causing microbes and stimulate enrichment of beneficial microbe that have potential to promote crop health [43]. In the present study, bacterial richness and diversity of each rhizospheric soil sample was measured and we observed that samples from Sheikhupura soil contain a higher bacterial richness (Chao1) and diversity (Shannon) than the other investigated sites. Altogether, this was found to contain a relatively high available P, available NO3 and low EC as compared to the other investigated sites of Basmati rice growing Kalar belt. It appeared that these soil parameters may contribute to shape the community structure of diverse microflora. It has been previously reported that soil properties directly influence the diversity of bacteria [44,45].

The understanding and exploitation of phyllosphere community in addition to rhizosphere would be helpful to suppress the leaf inhabiting pathogen and promote the crop health. According to best of our knowledge, no study is available for comparison of bacterial community in phyllosphere and rhizosphere of rice variety “Super Basmati”. In the present study, rhizosphere soil microflora was found to be very diverse in its bacterial composition with 208 distinct OTUs in accordance with the known complexity of the system as compared to phyllosphere. Previous scientists also observed higher population of bacteria in rhizospheric soil [46]. In present findings, we observed that sequences related to few genera detected in rhizosphere were not detected in the phyllosphere and similarly few genera were also specific for phyllosphere environment. These results indicated that bacteria belonging to these genera may have potential to survive in diverse environmental conditions. We detected 15 genera in the both compartment, *GpI, Bradyrhizobium, Microvirga, Sphingomonas, Brevundimonas, Pseudomonas, Thiobacillus, Ramlibacter, Syntrophobacter, Bacillus, Hydrogenispora, Sporacetigenium, Turicibacter, Gp6, Flavobacterium*. Here *Pseudomonas* and *Bacillus* is well known PGPR but *Pseudomonas* belong to biosafety level 2. We may targe other bacteria belonging to biosafetety level 1 for foliar as well as soil/seed inoculation for enhanced gowth of Basmati rice.

In the present study, we compared the bacterial diversity from different rice growing areas. On the basis of diversity indices, we found bacterial population in Sheikhupura and Gujranwala (areas of Kalar belt) are more diverse as compared to NIBGE field, Faisalabad. Moreover, large number of sequences 79-82% were unclassified at genus level in these sites. Bacterial population of Faisalabad site is significantly different from Sheikhupura and Gujranwala as 50 genera were specifically detected in Faisalabad. Comparison of bacterial diversity inhabiting the phyllosphere and rhizosphere of rice revealed that bacterial population associated with phyllosphere is less diverse as compared to rhizosphere. Few common persistent genera were detected in both habitat showed that these genera can tolerate nutrient deficiency and adverse environment, that can be further used for development of cost-effective, environmental friendly bioinoculum. Bacterial diversity of rhizosphere soil in different sites is also important for application of compatible strain inoculum.

## Acknowledgment

Technical assistance of Zahid Iqbal Sajid, Muhammad Sarwar, Asghar, Imran and Zakir during lab and fieldwork is thankfully acknowledged.

